# Behavioral and neuronal dynamics in a rat model of obsessive-compulsive disorder

**DOI:** 10.64898/2026.06.15.730877

**Authors:** Adam-František Hanzlík, Ewa Szczurowska, Tereza Rydzyková, Eduard Kelemen

## Abstract

While cognitive and behavioral manifestations of obsessive-compulsive disorder (OCD) are well known, the neuronal dynamics underlying these symptoms remain poorly understood. Theoretical work suggests that changes in the attractor dynamics of neuronal networks towards increased stability and decreased flexibility might cause behavioral and cognitive symptoms of OCD. We used chronic treatment with the D2 and D3 dopamine receptor agonist quinpirole as a rat model of the disease. In this model, we examined changes in behavioral dynamics and, in a parallel experiment, changes in organization of neuronal activity in the hippocampus and anterior cingulate cortex (ACC). At the behavioral level, we observed increased locomotion and repetitive stereotypical trajectories in quinpirole-treated rats, with frequency of repetitions increasing over the course of the session. At the level of neuronal activity, a gradual increase in the firing rate of ACC neurons within a session paralleled the dynamics of behavioral stereotypy after quinpirole treatment. In quinpirole-treated rats, we observed increased stability in the temporal organization of hippocampal neuronal firing, but no increase in the stability of the spatial organization of discharge. The increased stability of hippocampal firing was observed at both the level of single neurons and coordinated activity of neuronal pairs, and was connected to modulation of activity by theta rhythm. Studying neuronal activity changes underlying behavioral and cognitive manifestations of brain disorders is crucial for understanding and treating brain pathologies.

**HIGHLIGHTS:**

- Dynamics of behavior and neuronal activity was characterized in rats after chronic quinpirole treatment, which is considered a model of obsessive-compulsive disorder.
- The quinpirole treatment led to repetitive stereotypical trajectories, with increasing frequency of repetitions over the course of a session.
- The quinpirole treatment was associated with more stable theta modulation of single-cell hippocampal firing within experimental sessions.
- Quinpirole increased stability in cell-pair correlations of hippocampal units within and between sessions.
- Quinpirole led to more stable coordination of local field potential activity between the hippocampus and anterior cingulate cortex at theta frequencies.

## INTRODUCTION

Obsessive-compulsive disorder (OCD) is a brain disorder with well-characterized cognitive and behavioral manifestations, but poorly understood underlying neuronal pathological mechanisms. In OCD, common behaviors such as washing, cleaning, or checking are organized with different, pathological dynamics. The behaviors themselves are not pathological, but their organization in time is; with emerging repetitive patterns, and overall loss of flexibility among the most conspicuous changes (Stein et al., 2019). In humans, these alternations in organization of behavior are accompanied by changes in dynamics of thought processes. Again, thoughts of concerns for safety and cleanliness are common in most people, but they manifest differently in OCD patients, were these thoughts become organized into repetitive patterns or loops, they lose their flexibility and are hard to escape or terminate (Gruner and Pittenger, 2017). According to an accepted view, the processes of thought and cognition that control behavior emerge from neuronal activity. Therefore, it is crucial to study the changes in the dynamics of neuronal activity that underlie the cognitive and behavioral dynamics in OCD.

Changes in the dopaminergic neurotransmitter system are implicated in OCD (Mota et al., 2026). In this study, we use a rat pharmacological model of the disease introduced by Szechtman et al. (1998), which is based on chronic exposure to a D2 and D3 receptor agonist quinpirole. This treatment leads to characteristic changes in rat’s behavior that resemble compulsive checking, such as repeated visits to some of the objects in an environment and ritual-like acts at locations of these objects. These manifestations have been considered and used as a relevant model of human OCD-like symptoms (Szechtman et al., 1998; De Haas et al., 2011; Stuchlik et al., 2016).

In the first part of our study, we characterize the dynamic organization of rat behavior in space and time in the quinpirole model. In extension of previous observations (Szechtman et al., 1998; De Haas et al., 2011; Stuchlik et al., 2016), we hypothesize that after quinpirole treatment, the behavior will be less variable, more predictable, and organized into repeating patterns and sequences. In the second part of the work, we explore the dynamic organization of neuronal activity in the same model. Theoretical work (Rolls et al., 2008; Rolls, 2012) has proposed that OCD symptoms are associated with altered neuronal attractor dynamics with deeper attractor states, which may manifest in more stably organized neuronal activity. We assess the dynamics of organization of neuronal activity in space and time, the stability of organization of single cell activity, and the coordination of activity among neurons in neuronal groups.

We focus on neuronal activity in two cortical structures implicated in the disease: the anterior cingulate cortex (ACC) and the hippocampus. ACC is part of one of the cortico-striato-thalamo-cortical loops – neuronal circuits involved in OCD (Posner et al., 2014). Evidence implicating ACC in OCD includes observations that OCD patients perform differently in many cognitive tasks that rely on the ACC, such as performance monitoring and error processing (Brown and Braver, 2005; Li et al., 2019; Van De Veerdonk et al., 2023). OCD has been associated with increased electrophysiological and blood oxygenation measurements in ACC (Perani et al., 1995). ACC was overactive in OCD patients undergoing a fast reaction time task, in which they had to suppress a compulsion to respond to conflicting stimuli (Maltby et al., 2005). Simultaneous EEG and fMRI recordings showed a correlation between event-related potentials and BOLD signal from the subgenual ACC in OCD patients undergoing an error-processing task (Grützmann et al., 2016). LORETA analysis of sources of cranial EEG in OCD patients revealed increased power of slow oscillations (2-6 Hz) in subgenual part of the ACC (Kopřivová et al., 2011).

There is growing evidence for anatomical alterations of the hippocampal formation in OCD patients. Decrease in hippocampal volume (Boedhoe et al., 2017; Honda et al., 2017) and other anatomical anomalies of the hippocampal formation (Hong et al., 2007) were observed in the disease. Lower hippocampal volume was associated with a cluster of symptoms characterized by high checking and ordering behavior (Reess et al., 2017). There is evidence that hippocampal plasticity is altered in an animal model of OCD; decreased expression of plasticity-related immediate early genes was found in the hippocampal CA1 area of rats with pharmacologically induced OCD-like behavior (Brozka et al., 2021). The hippocampus and ACC are thus relevant targets for exploration of neuronal mechanisms involved in OCD. In this work, we will characterize the dynamic structure of behavior and neuronal activity in a rat model of OCD, focusing on the stability in observed behavioral and neuronal activity patterns.

## METHODS

### Animals

Adult male rats of Long-Evans strain (Velaz s.r.o., Prague, Czechia) were used in the study. Seventeen rats were used in Experiment 1 (behavioral study), and ten rats were used in Experiment 2 (electrophysiology study). The rats were housed in an animal facility with controlled temperature (22±2 °C), humidity (50±20%), and12/12-hour light/dark cycle (lights on at 6:00 a.m.). During experiments, the food was restricted so that animals remained at >80% of their expected free-feeding weight. All animal experiments were approved by the Ministry of Health of the Czech Republic (project number 42/2020) and performed in accordance with the Animal Protection Act of the Czech Republic and the European Union directive 2010/63/EC.

### Pharmacological manipulations

Pharmacological manipulation followed the quinpirole sensitization protocol described earlier (Szechtman et al., 1998). Animals in the quinpirole-treated group (QUINP) received repeated subcutaneous injections of quinpirole (0.5 mg per kg of body weight) administered three times per week, up to 10 injections. A solution of 0.5mg/mL quinpirole in sterile saline was used (Bio-Techne R&D Systems, Prague, Czechia). Animals of the control group (CONTR) received saline injections of equal volume with the same schedule.

### Behavioral protocol

One hour after each injection, animals were placed for 30 minutes in a square arena (86 × 86 cm, wall height 60 cm) containing three objects: a small ceramic flowerpot, a round plastic cup, and a square enamel mug. This setup was later used as the familiar configuration (FAM; Fig. 2A). The arena, objects, and their positions remained consistent across habituation sessions. Food pellets were provided during arena exploration. In both Experiment 1 (behavioral study) and Experiment 2 (electrophysiological study), data were acquired during two consecutive sessions. The first session was conducted in the familiar object configuration (FAM), and the second in a novel configuration (NOV), in which the positions of the objects were changed (Fig. 3A). Behavioral data in Experiment 1 were collected 60 min after the last quinpirole injection. Electrophysiological data in Experiment 2 were collected up to 10 days after the last injection.

**Figure 1.**
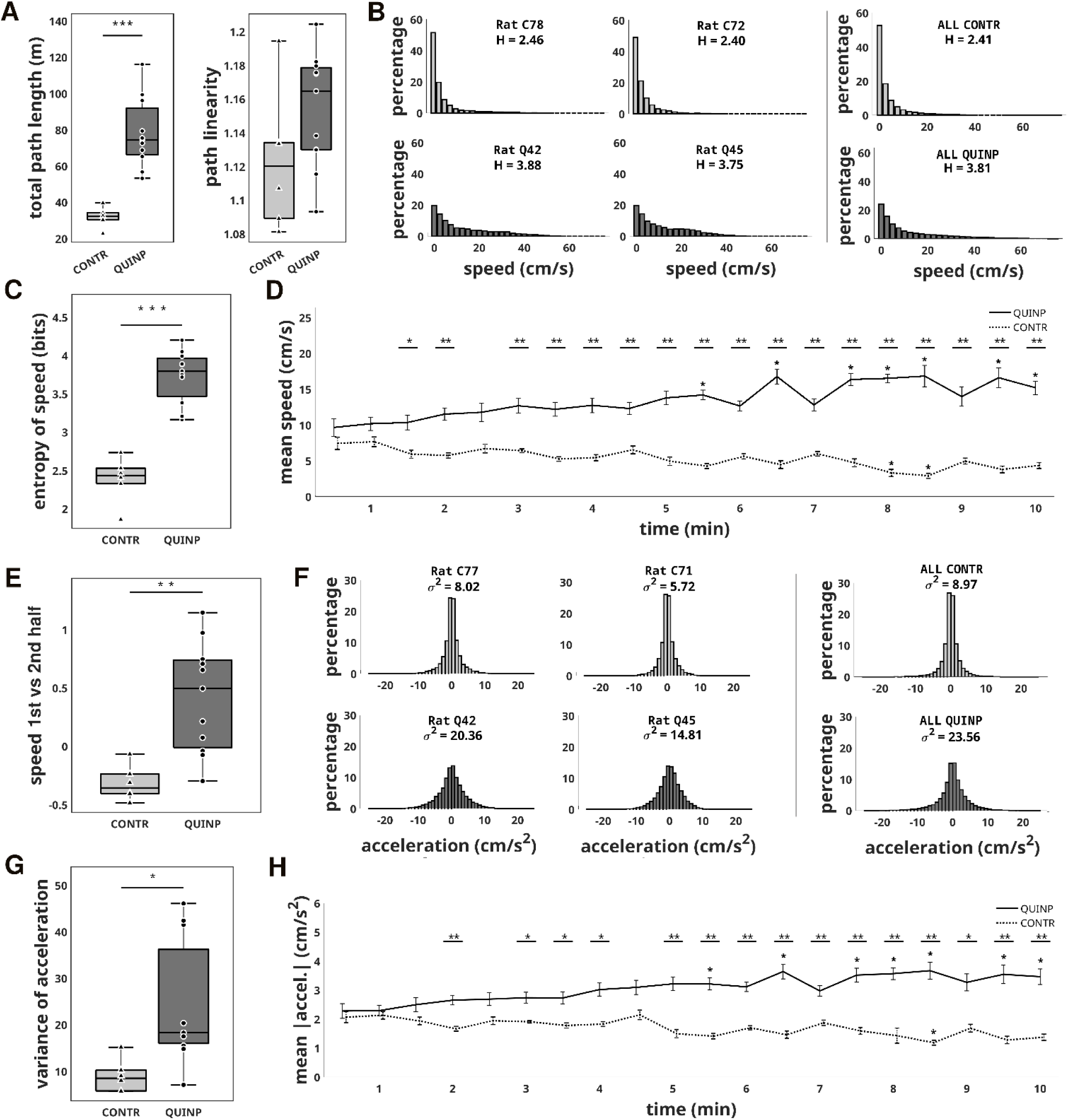
Locomotor behavior – Experiment 1. **(A)** Total path length and path linearity. **(B)** Distributions of speed of locomotion. Left and middle panels: Examples from two CONTR rats (top) and two QUINP rats (bottom). Right panels: Average distribution of speed from the CONTR and QUINP groups. **(C)** Entropy of speed distributions. **(D)** Changes in speed within a session in the CONTR and QUINP groups. Asterisks in the top part of the chart indicate significant differences between groups. Asterisks directly above data points indicate a significant difference from the beginning of the session at t=30s. **(E)** The difference of median speed between the 2nd half and the 1st half of a session, divided by the median speed in the 1st half of the session. **(F)** Distributions of acceleration. Left and middle panels: Examples from two CONTR rats and two QUINP rats. Right panels: Average distribution of acceleration in the CONTR and QUINP group. **(G)** Variance of acceleration distributions shown in (F). **(H)** Mean absolute value of acceleration across 30s intervals in the CONTR and QUINP groups. Significance marked as in (D). (* p<0.05, ** p<0.01. *** p<0.001)

**Figure 2.**
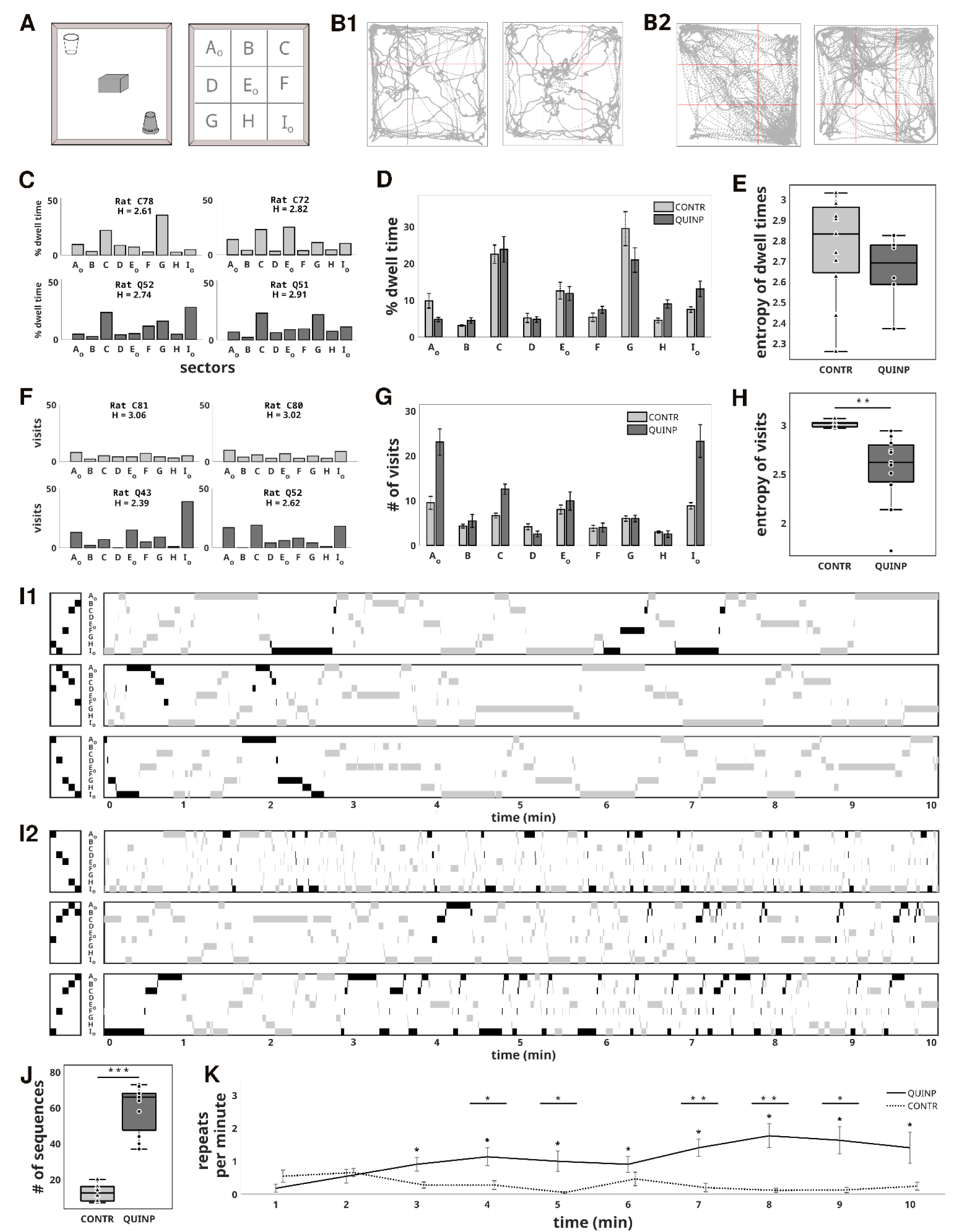
Spatial organization of rat movements on the arena – Experiment 1. **(A)** Drawing of the arena with three objects in the familiar configuration (FAM). For analysis, the arena was divided into nine sectors. **(B)** Examples of trajectories from two CONTR rats (B1) and two QUINP rats (B2). **(C)** Examples of relative sector dwell times from two sessions in CONTR (top) and two sessions in QUINP (bottom) rats. Entropy value H is printed above each distribution. **(D)** Average dwell times in different sectors for the CONTR and QUINP groups. **(E)** Average entropy of distributions of dwell times for CONTR and QUINP rats. **(F)** Examples showing the number of visits to individual sectors from two sessions in CONTR (top) and two sessions in QUINP (bottom) rats. **(G)** Average number of visits to different sectors for CONTR and QUINP rats. **(H)** Average entropies of distributions of visits for CONTR and QUINP rats. **(I)** Movements through the nine sectors within a single experimental session. Three sessions from CONTR rats (I1) and three sessions from QUINP rats (I2) are shown. Gray rectangles depict the time spent in the nine sectors. In each diagram, the most-repeated sequence of five subsequently visited sectors is highlighted in black. **(J)** The total count of five-sector sequences that repeated more than twice. **(K)** Number of repeats of five-sector sequences that repeated more than twice as a function of session time. Asterisks at the top indicate significant differences between groups. Asterisks directly above data points indicate a significant difference from the data at t=1 min.

**Figure 3.**
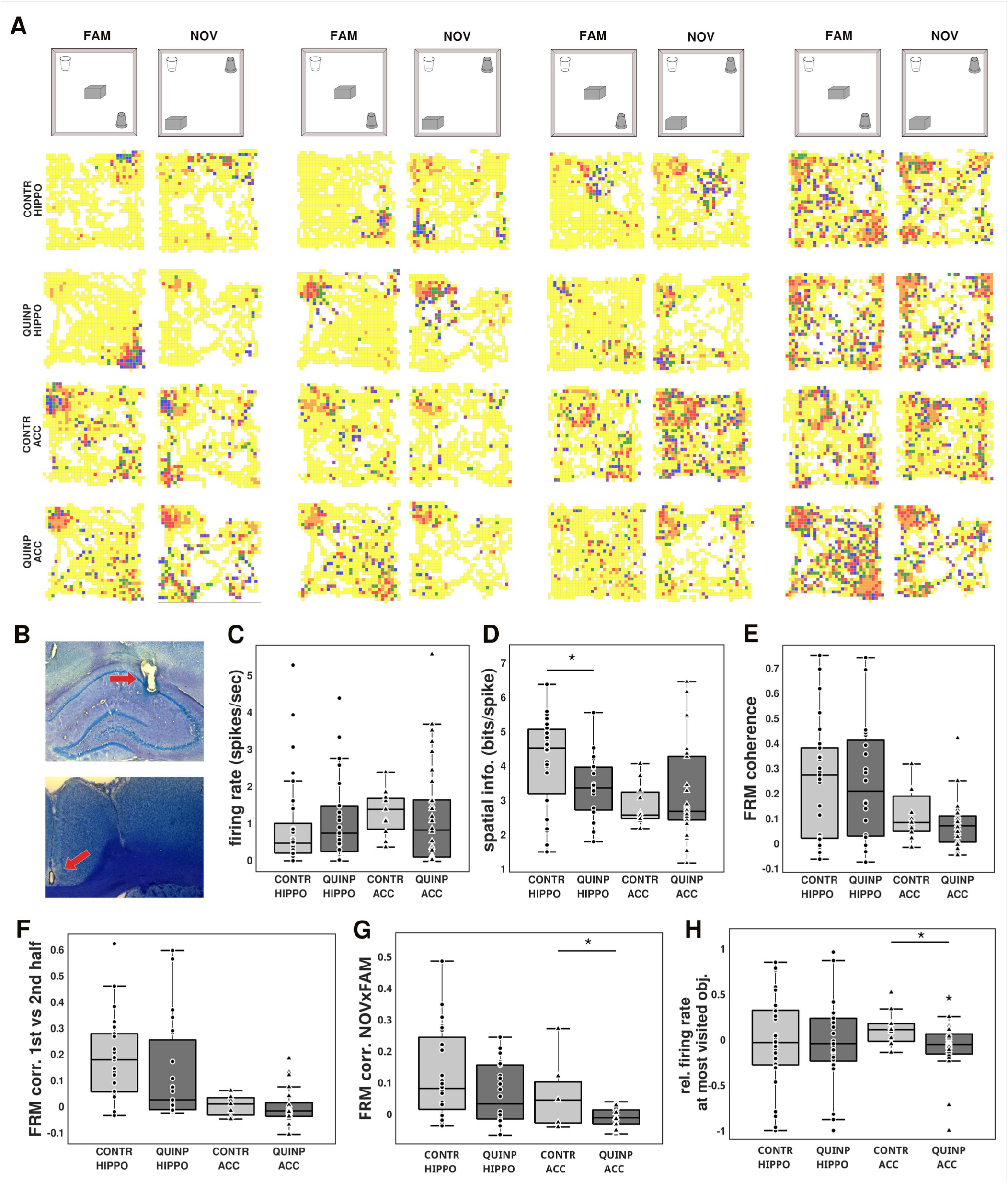
Spatial organization of neuronal activity – Experiment 2. **(A)** Examples of firing rate maps of the hippocampal and ACC units from QUINP and CONTR animals. Each unit’s activity is shown in sessions with the familiar (FAM) and novel (NOV) object configurations. Darker and colder colors indicate locations with higher firing rates; brighter and warmer colors indicate lower firing rates; yellow means a firing rate of 0 spikes per second. **(B)** Examples of histological tracking of electrode positions in the hippocampus (top) and in the ACC (bottom). **(C)** Average firing rate. **(D)** Average spatial information. **(E)** Average firing rate map coherence. **(F)** Correlations between firing rate maps from the first and the second half of the FAM session. **(G)** Correlations between firing rate maps from FAM and NOV sessions. **(H)** Relative firing rate at the most visited object. Values > 0 indicate a higher firing rate near the most visited object compared to the mean firing rate in the remaining sectors. An asterisk directly above a boxplot marks a significant difference from zero.

### Surgical and electrophysiological methods

Surgical and electrophysiological methods used in Experiment 2 followed previously used protocols (Kelemen and Fenton, 2010; Ahuja et al., 2020). Electrodes made from Teflon-insulated nichrome wire (25 micrometers in diameter) were arranged in the “tetrode” configuration. Each rat was chronically implanted with four tetrodes (i.e., 16 electrode wires) in the dorsal CA1 region of the hippocampus and four tetrodes in the ipsilateral ACC. The electrodes were built into a “VersaDrive” implant that allowed positioning of electrodes in the vertical direction (Axona Ltd, St. Albans, UK). Surgery was performed under isoflurane anesthesia (dose for induction: 5%, dose for maintenance: 1.5-3%) in close to aseptic conditions. A rat was placed into a stereotaxic apparatus (Neurostar, Tübingen, Germany), the skin on the skull was opened, and the skull was cleaned. Four holes were drilled for anchoring bone screws, and two holes served for the insertion of intracerebral electrodes. The hippocampal electrodes were aimed 3.8 mm posterior, 2.5 mm lateral from Bregma, and initially positioned 1.5 mm below the brain surface. ACC electrodes were aimed 1.5 mm anterior, 0.5 mm lateral from Bregma, and 0.5 mm below the brain surface. Two of the bone screws in the occipital bone served as grounding and reference electrodes. Surgeries were performed within the pharmacological sensitization protocol, between the 6^th^ and 9^th^ injections of either quinpirole or saline.

During electrophysiological recordings, the rats’ implants were connected to head-stage preamplifiers and, via a cable, to an amplifier and a recording system (Axona Ltd, St. Albans, UK). Local field potential (LFP) signals were low-pass filtered at 100 Hz, digitized, and stored at 250 Hz. Single-unit signals were filtered between 300 Hz and 7 kHz and digitized and stored at 48 kHz and 24 bit. Prior to the recording session, the tetrodes were gradually lowered to the desired depth in a series of screening sessions. After the experiment concluded, the animals were sacrificed by isoflurane overdose and subsequent decapitation. The position of tetrodes was marked by passing 15µA current through each electrode. The brains were extracted and infused by 8% paraformaldehyde, followed by 30% sucrose, then frozen, cut into 50 µm sections, and stained with Toluidine blue. The position of tetrodes was verified using optical microscopy (Fig. 3B).

### Analysis of behavior

Each experimental session was video recorded, the rat’s position in the arena was sampled at 30 Hz and tracked using Ethovision XT software (Noldus, Wageningen, the Netherlands). To analyze basic locomotor activity, instantaneous speed and acceleration were calculated over 267ms intervals (eight position frames). Linearity of locomotion was calculated as the ratio of the observed length of the trajectory and the shortest possible trajectory between two positions over 1-second intervals. To analyze the spatial organization of behavior, the surface of the arena was divided into nine square sectors (Fig. 2A) and distributions of dwell time and number of visits in the nine sectors were calculated (Fig. 2). The homogeneity of a distribution was assessed by entropy using the following formula:

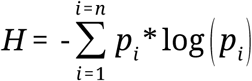

where *H* is entropy, *n* is the number of measurement categories (nine sectors of the arena), *p_i_* is the probability of the i^th^ category, and *log* is the binary logarithm. Higher entropy is characteristic of more homogenous distributions; lower entropy reflects less homogeneity.

### Analysis of neuronal activity

Action potentials of individual neurons were sorted using KlustaKwik v.3 (Kadir et al., 2014) and then manually curated using TINT cluster-cutting software (Axona, St. Albans, UK). Putative interneurons were differentiated from putative principal cells based on standard criteria (Fox and Ranck, 1981; Csicsvari et al., 1999).

To study the organization of a unit’s discharge in space, we constructed firing rate maps (FRM) (Muller et al., 1987). The spatial organization of firing was quantified by FRM coherence (Muller and Kubie, 1989) and spatial information content (Skaggs et al., 1992). To investigate the stability of spatial organization within a session, we calculated Pearson’s correlation of FRMs from the first and the second half of a session. We used raw FRMs for illustrations and for calculating coherence and information content. All other analyses of FRMs, such as correlations between halves of a session or correlations between FAM and NOV sessions, used FRMs smoothened using a Gaussian kernel with a standard deviation of 0.5. To assess a unit’s preference to discharge near the most-visited object, we calculated the normalized firing rate index using the formula:

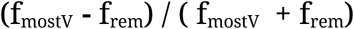

where f_mostV_ is the firing rate in the sector with the most-visited object and f_rem_ is the mean firing rate in the remaining sectors. Only units with an average firing rate > 0.1 Hz were included in the analyses of the spatial organization of firing.

To investigate the temporal structure in the firing of individual units and to assess theta modulation and bursting we calculated autocorrelation functions of spike trains. The *theta index* was obtained from the Fast Fourier Transform (FFT) of an autocorrelogram by dividing the total power in the theta frequency range (5-10 Hz) by the total power between 2-50 Hz. *Bursting score* was defined as the probability that the inter-spike interval of a spike train is less than 10 ms.

We calculated cross-correlations to study co-activation between pairs of units. The cross-correlation theta index was calculated in a way analogous to the autocorrelation theta index. We also compared the value of cross-correlation function at ±30 ms around zero relative to other bins by the *near-zero index*:

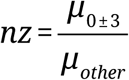

where μ_0±3_ is the mean of the cross-correlation function over si× 10 ms bins around zero, and μ_other_ is the mean of the cross-correlation function over all other bins. The *symmetry score* of a cross-correlation was calculated as

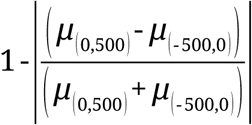

where μ_(0,500)_ is the mean of the cross-correlation function from 0 to 500 ms, and μ_(−500,0)_ is the mean of the cross-correlation function from −500 to 0 ms.

To further analyze correlations of activity between neuronal pairs, the recording session was divided into one-second-long intervals, and each interval was subdivided into 20-ms-long bins. Only bins with non-zero spike counts were used. For each one-second interval, Pearson’s correlation coefficient between spike counts within the bins was computed. Pearson’s coefficients for subsequent one-second intervals were compared to assess the stability of pairwise neuronal coactivation within the experimental session. Other interval lengths and bin sizes were also analyzed.

The LFP signal was divided into two-second intervals, and for each interval FFT was applied. For the analysis of theta phase difference and phase locking, the LFP signal was filtered between 5-12 Hz using a zero-phase Chebyshev type II filter, and Hilbert transformed to obtain the amplitude and phase. To assess theta synchronization between ACC and hippocampus, time series of momentary phase differences between hippocampal and ACC theta were calculated (Fig. 6A, B). To calculate theta phase locking of unit firing, the distribution of the unit firing rate as a function of theta phase was calculated (Fig. 6G, H, I, J), and the mean phase and mean modulation strength of theta phase locking were determined. LFP traces from the respective brain regions (i.e., hippocampal recording for hippocampal units and ACC recording for ACC units) were used for phase locking analysis.

### Statistics

MATLAB software (Mathworks, Natick, MA, USA) was used for statistical comparison. Where appropriate, we used Student’s t-test, ANOVA, and Newman-Keuls post hoc test. When the condition of normality was not met, we performed Wilcoxon rank-sum tests for two independent variables and the Wilcoxon signed-rank test for paired variables. A two-sample Kolmogorov-Smirnov test was performed to compare the overall shape of two distributions. Wald-Wolfowitz runs test was used to test the randomness of data sequence. All Pearson’s correlation values and FRM coherence values were Fisher-transformed before statistical analysis.

## RESULTS

### Behavior (Experiment 1)

We assessed the behavior of 11 quinpirole-treated (QUINP) rats and 6 control (CONTR) rats. Rats in the QUINP group walked longer total distances than those in the CONTR group (t(15)=5.73, p<0.001; Fig. 1A). There was no significant difference in linearity between the two groups (t(15)=1.63, p=0.12; Fig. 1A), so the difference in path length was not the result of a more meandering walking pattern in the QUINP group. The longer total paths in QUINP rats imply a higher average speed of locomotion. Both groups spent a large proportion of time moving at low speeds (<4 cm/s). High-speed intervals were less frequent, but they were more often observed in the QUINP group than in CONTR rats (Fig. 1B). Homogeneity of speed distributions in each rat was quantified by entropy, which was significantly higher in QUINP compared to CONTR rats (t(15)=8.14, p<0.001; Fig. 1C). While the speed of CONTR rats was not changing much within a session, or was getting slightly slower with time, in QUINP rats there was a gradual increase in speed over the course of the session (t(15)=3.73, p<0.05 for 1^st^ vs 2^nd^ half; Fig. 1D and Fig. 1E). Acceleration was assessed next (Fig. 1F). Intervals with higher absolute value of accelerations were more prominent in the QUINP group, reflecting faster, more abrupt changes in speed (t(15)=-2.67, p<0.05 for variance of acceleration; Fig. 1F and 1G). Entropy of distribution of accelerations was significantly higher in QUINP group (t(15)=-5.92, p<0.001; data not shown). There was a gradual increase in the absolute value of acceleration within a session in the QUINP group and no change or slight decrease in the CONTR group, similar to the trend in speed (t(15)=-2.77; p<0.05 for 1^st^ vs 2^nd^ half; Fig. 1H).

Trajectories in QUINP rats were more regular, while the CONTR group displayed more random walking patterns (Fig. 2B1 and 2B2). For further quantitative analysis, the arena was divided into nine square sectors (Fig. 2A), and the dwell time (the total time spent) in each sector was determined. Two-way ANOVA indicated significant effect of sectors (F(8,135)=22.95, p<0.001), no significant interaction (F(8,135)=1.84, p=0.07; Fig. 2C and 2D). Post hoc comparisons revealed that rats spent significantly more time in the corners of the arena that did not contain objects (sectors ‘C’ and ‘G’; Fig. 2D). The homogeneity of dwell times for each rat was quantified by entropy and there were no significant differences between QUINP and CONTR groups (t(15)=-0.95; p=0.36; Fig. 2E).

For the next set of behavioral analyses, we focused on spatial patterns of the rats’ movements in the arena. We identified visits to the nine sectors that were at least two seconds long, and we ignored shorter passes through sectors in this analysis. The most visited sectors in the QUINP group were the corners of the arena that contained objects (sectors ‘A’ and ‘I’). Average entropy of visit distributions was lower in the QUINP group compared to the CONTR group (t(15)=-3.08, p<0.01; Fig. 2H). This indicates that QUINP rats had a stronger preference for their most visited sectors than CONTR rats had for their most visited sectors (Fig. 2F).

Next, we analyzed stereotypy in sequences of visits to different sectors. For each recording session, we identified sequences of subsequently visited sectors. Examples of such sequences of five subsequently visited sectors in three CONTR rats and three QUINP rats are shown in Figure 2I. We observed many more repeats of the same sequences in QUINP rats compared to CONTR rats. For 5-sector sequences, QUINP rats showed higher counts of sequences (t(15)=8.36, p<0.001) and more repetitions of sequences (ranksum = 250543.3, p<0.001, Wilcoxon rank-sum test; Fig. 2J). The trend was similar for three-, four-, and six-sector sequences (data not shown). The frequency of repeats in QUINP rats increased later in the recording session (Fig. 2K).

### Neuronal activity (Experiment 2)

Unit activity was recorded from the hippocampus and ACC of quinpirole-treated (QUINP, N=4) and control (CONTR, N=5) rats in sessions with familiar object configuration (FAM) followed by a session with novel object configuration (NOV). We assessed the organization of neuronal firing and its stability in the domains of space and time.

### Organization of neuronal activity in space

There was no significant effect of treatment on the average firing rate of hippocampal (ranksum=748, p=0.40, Wilcoxon rank-sum test) or ACC units (ranksum=296, p=0.26; Fig. 3C). Examples of FRMs of hippocampal and ACC neurons are shown in Figure 3A. The quality of spatial organization of principal cell firing was quantified by rate map coherence (Muller et al., 1987) and spatial information (Skaggs et al., 1992). There was no significant effect of treatment on the coherence of FRMs of hippocampal units (t(45)=0.19, p=0.85) or ACC units (t(35)=1.02, p=0.31; Fig. 3E). Spatial information was significantly lower in hippocampal units from QUINP rats than in CONTR rats (t(45)=2.34, p=0.02), there was no such difference in the ACC neurons (t(35)=-0.77, p=0.45; Fig. 3D). The stability of spatial organization of firing within a recording session was assessed by comparing FRMs constructed for the first half and the second half of each session. We observed no effect of quinpirole treatment on the correlations of FRMs between the first and the second half of a session in hippocampal (t(46)=1.30, p=0.20) or ACC units (t(26)=0.03, p=0.97; Fig. 3F).

Next, we assessed the stability of the spatial organization of neuronal firing in response to environmental changes between FAM and NOV sessions. The tendency of hippocampal FRMs from QUINP rats to be less correlated than hippocampal FRMs from CONTR rats was not significant (t(44)=1.77, p=0.08; Fig. 3G). The maps of ACC units were significantly less correlated in QUINP group than in CONTR rats (t(24)=2.43, p=0.02; Fig. 3G).

Because behavioral analysis presented earlier revealed an increased number of visits to particular squares with objects in the QUINP group, we next compared the firing rate of units in a sector containing the most visited object with the mean firing rate in the remaining eight sectors (Fig. 3H). Hippocampal units’ average firing rate was not affected by the most-visited object, and there was no difference in tendency to fire near the most visited object (expressed as normalized firing rate index – see Methods) between QUINP and CONTR rats (t(52)=0.25, p=0.80). The ACC units in QUINP rats discharged near the most-visited object less than in the other sectors (t(33)=-2.31, p=0.03), and their tendency to fire near the most-visited object (normalized firing rate index) was significantly lower when compared to ACC units from CONTR rats (t(43)=2.32, p=0.03).

### Organization of single-unit activity in time

Next, we analyzed the organization of neuronal firing in time and its stability. First, we inspected the change of the firing rate of putative principal cells within the course of a recording session (Fig. 4A and 4B). There was no significant change in firing rates of hippocampal units within a session (Wald-Wolfowitz runs test, Z=-0.05, p=1 and Z=0.28, p=0.80 for QUINP and CONTR groups, respectively; Fig. 4A). ACC units from QUINP rats increased their firing rate approximately half-way through the FAM session (Z=-2.10, p<0.05) whereas ACC units from CONTR rats did not (Z=-0.23, p=0.83; Fig. 4B).

**Figure 4.**
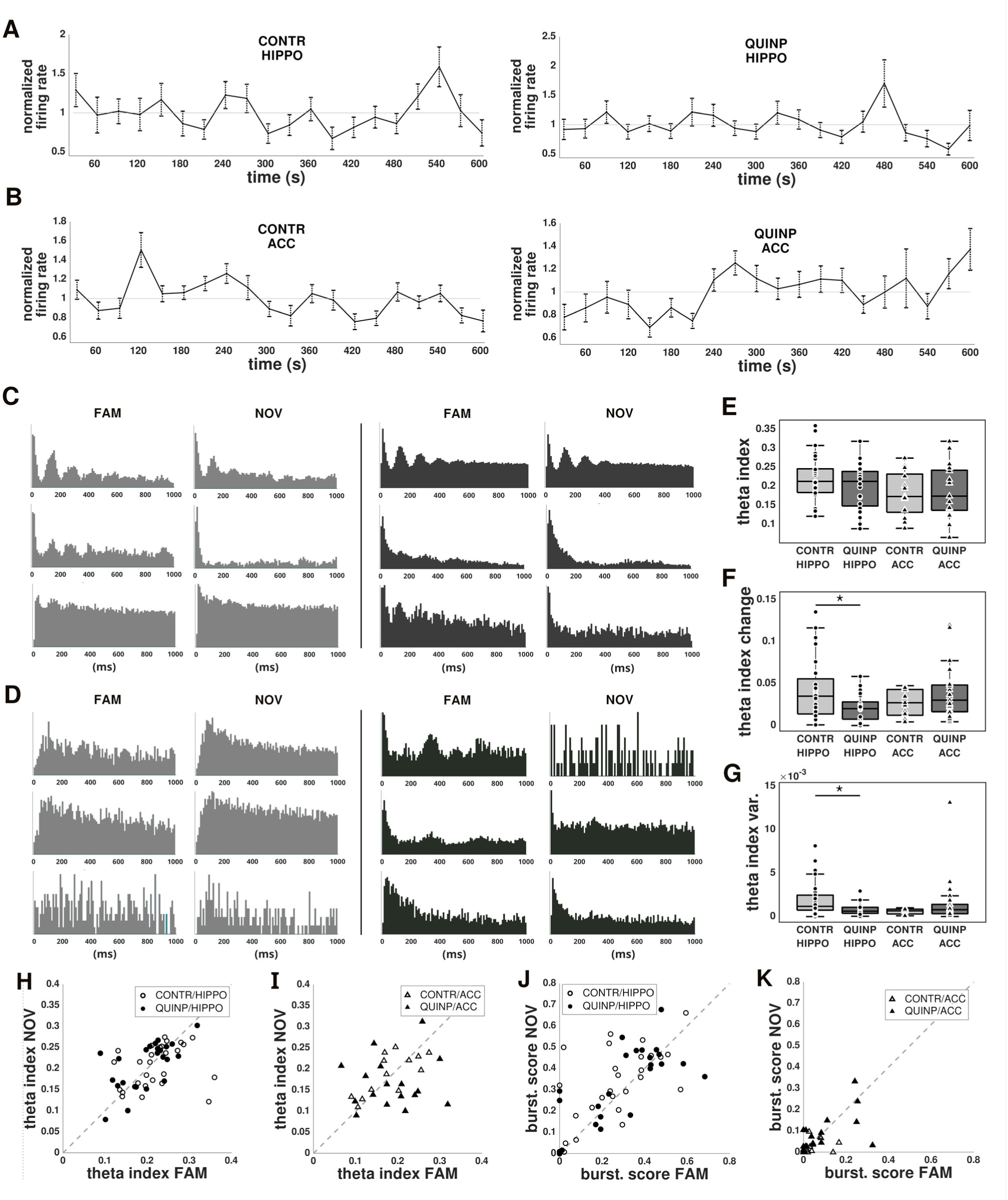
Organization of activity of single neurons in time – Experiment 2. **(A)** and **(B)** Changes in firing rate within a recording session for hippocampal units (A) and ACC units (B) in the CONTR (left) and QUINP (right) group. **(C)** and **(D)** Examples of autocorrelograms of hippocampal (C) and ACC (D) units from CONTR (gray) and QUINP (black) rats. The same units are shown in the FAM and NOV recording configuration. **(E)** Mean values of the autocorrelation theta index. **(F)** The change in theta index between the first and second half of the FAM session. **(G)** Variance of theta index across 100-s intervals. **(H)** and **(I)** Theta index in FAM and NOV sessions for hippocampal (H) and ACC (I) units. **(J)** and **(K)** Bursting score in FAM and NOV sessions for hippocampal (J) and ACC (K) units.

Autocorrelations (Fig. 4C and 4D) reveal characteristic features in the organization of single neuron discharge in time, such as potential modulation of neuronal activity by theta rhythm and bursting in neuronal firing. Quinpirole treatment had no effect on the average theta index (t(49)=1.36, p=0.18 for hippocampal and t(28)=-0.23, p=0.82 for ACC units; Fig. 4E), and the average bursting score also did not significantly differ between treatment groups (t(49)=-0.64, p=0.52 for hippocampal and t(28),p=0.82 for ACC units). We studied the stability of these characteristics between the first and the second half of the FAM recording session. For hippocampal units, the theta index changed significantly less in QUINP rats than in the CONTR group (t(46)=2.41, p<0.05; Fig 4F), suggesting increased stability after quinpirole treatment. Significant difference of intra-session change was not found in units from the ACC (t(36)=-1.23, p=0.23). To characterize the stability of firing on shorter time scales, each session was divided into 100-second-long intervals, and the theta index was computed for each interval. For hippocampal units, the intra-session variance of the theta index was significantly lower in QUINP rats than in the CONTR group (t(46)=2.31, p=0.03; Fig. 4G). No significant effect of treatment on the intra-session variance of the theta index was found in units from the ACC (t(36)=-1.16, p=0.25). The rate of change of the theta index between subsequent 100-sec intervals was also lower in QUINP than in the CONTR group for hippocampal units (t(43)=3.00, p<0.01), but not for ACC units (t(35)=-1.61, p=0.12). Dividing the session into shorter (60 seconds) or longer (120 seconds) intervals yielded similar results. Quinpirole had no significant effect on the change of bursting score between the first and second half of the FAM session (ranksum=603, p=0.64 for hippocampal and ranksum=239, p=0.54 for ACC units; Wilcoxon rank-sum test).

Next, we evaluated the stability of features of autocorrelations between the FAM and NOV session. We calculated correlations of parameters for the same units in the FAM and in the NOV sessions, and we also quantified the inter-session change by taking the absolute value of the difference of parameters in the FAM and NOV sessions. There was a significant positive correlation of theta index between NOV and FAM sessions in hippocampal units from QUINP rats (r=0.64, p<0.001, Fig. 4H) but not in hippocampal units from CONTR rats (r=0.25, p=0.20). There was no significant effect of treatment on the between-session change of theta index in the hippocampus (ranksum=706, p=0.95, Wilcoxon rank-sum test). For ACC units, CONTR rats displayed a significant correlation of theta index (r=0.72, p<0.01, Fig. 4I), QUINP rats did not (r=0.09, p=0.75). Theta index changed significantly more between sessions in the ACC units from QUINP rats (ranksum=148, p<0.05, Wilcoxon rank-sum test).

There was a significant positive correlation between bursting scores in the FAM and NOV sessions for hippocampal units from QUINP rats (r=0.74, p<0.001) as well as from CONTR rats (r=0.62, p<0.001; Fig. 4J). For ACC units, only the QUINP group showed a significant correlation of bursting scores between the FAM and NOV sessions (r=0.61, p<0.01; Fig. 4K). There was no significant difference between QUINP and CONTR groups in how the bursting score changed between sessions, neither in the hippocampus (ranksum=710.2, p=0.88) nor in the ACC (ranksum=166.5, p=0.15, Wilcoxon rank-sum tests).

### Coordination of neuronal activity between neuronal pairs

To analyze the coordination of activity between pairs of neurons and its stability, we first computed cross-correlations of simultaneously recorded units (Fig. 5A and 5B). Modulation of co-firing by theta rhythm was again assessed by theta index. Co-activation of neurons within a few milliseconds manifested as a peak near zero, and the tendency not to fire together manifested as through near zero. These tendencies were quantified by near-zero index. The tendency of one cell to activate before or after another was measured by the symmetry score. To assess within-session stability of cross-correlations, we compared cross-correlation parameters between the first and the second half of recording sessions. There was no effect of quinpirole on the theta index change between halves of the FAM session for neither hippocampal (t(183)=-1.30, p=0.20) nor ACC pairs (t(142)=-0.01, p=0.99; Fig. 5E). Of note, there was a significant difference between QUINP and CONTR groups in the average theta index of ACC pairs (t(168)=-2.56, p=0.01; Fig. 5C). Hippocampal pairs from QUINP rats changed their symmetry score significantly less than hippocampal pairs from CONTR animals (p<0.01, ranksum=20031, Wilcoxon rank-sum test; Fig. 5F). There was no significant treatment effect on the absolute value of near-zero index change between the first and the second half of the FAM session for neither hippocampal (ranksum=18929, p=0.41) nor for ACC (ranksum=1008, p=0.06) pairs.

**Figure 5.**
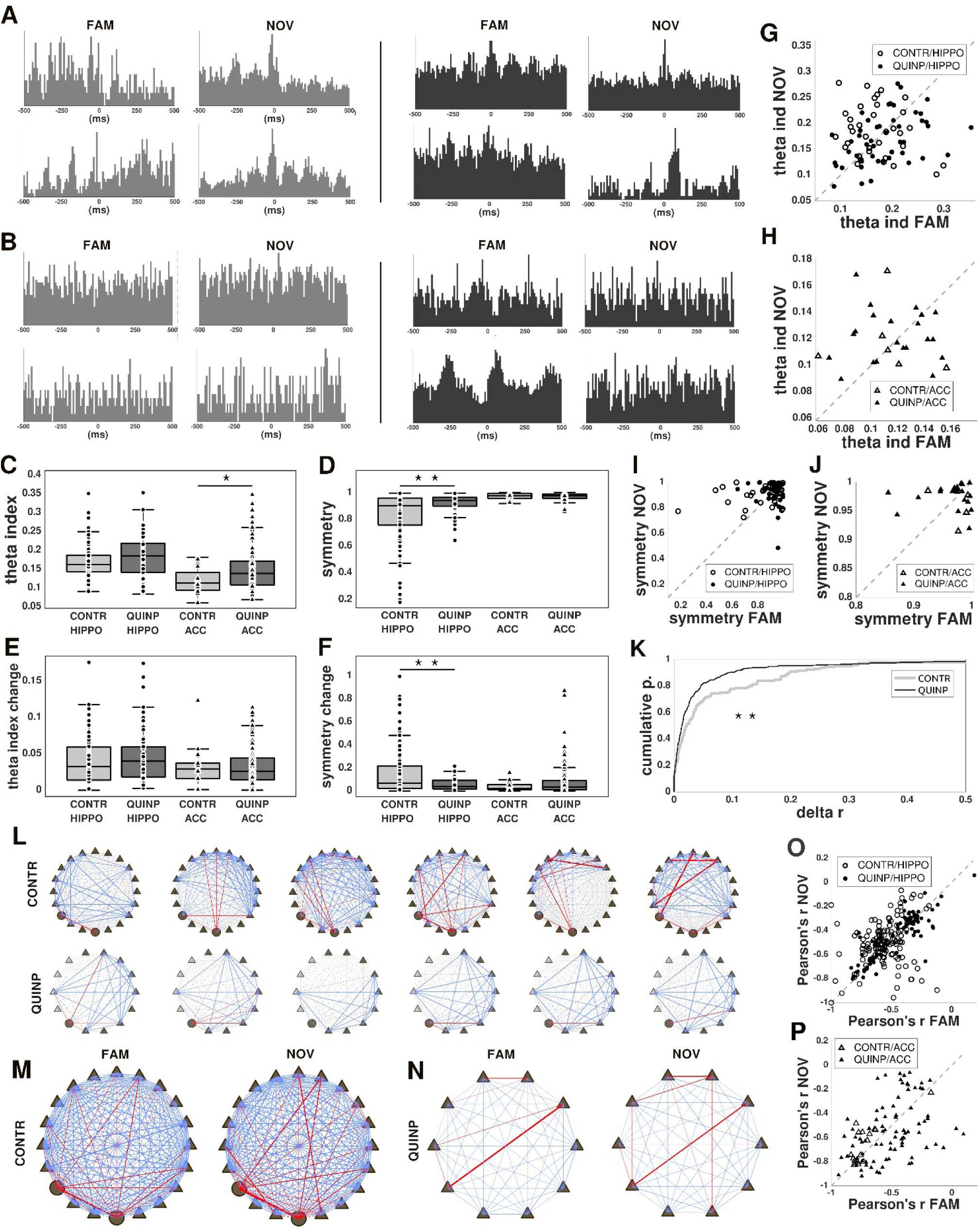
Organization of activity between neurons – Experiment 2. **(A)** and **(B)** Examples of cross-correlograms of pairs of hippocampal (A) and ACC (B) units from CONTR (gray) and QUINP (black) rats. **(C)** Mean theta index of cross-correlograms. **(D)** Mean symmetry score of cross-correlograms. **(E)** The change in theta index of cross-correlograms between the first and the second half of the FAM session. **(F)** The change in the symmetry of cross-correlograms between the first and the second half of the FAM session. **(G)** and **(H)** Theta index of cross-correlograms in the FAM and NOV session for hippocampal (G) and ACC (H) unit pairs. **(I)** and **(J)** Symmetry score of cross-correlograms in the FAM and NOV sessions for hippocampal (I) and ACC (J) unit pairs. **(K)** Cumulative distribution of the differences of Pearson’s correlation coefficients calculated between consecutive 1s bins (delta r) in the FAM session. Lower values of delta r indicate more stable correlation patterns. **(L), (M)**, and **(N)** Diagrams of Pearson’s correlation coefficients between pairs of simultaneously recorded units drawn as lines between principal cells (triangles) and interneurons (circles). Positive correlations are depicted as red lines, negative correlations as blue lines; the line’s thickness corresponds to the magnitude of correlation. Black fill denotes hippocampal, gray ACC units. **(L)** Examples of correlation diagrams from one CONTR (top row) and one QUINP (bottom row) animal. Each row consists of 6 consecutive five-second windows of neuronal activity. **(M)** and **(N)** Examples of correlation diagrams from one CONTR (M) and one QUINP (N) animal depicting the FAM (left) and NOV (right) session. Only hippocampal units are shown. **(O)** and **(P)** Pairwise correlation coefficients in FAM session plotted against coefficients from the same pairs in NOV session for hippocampal (O) and ACC (P) unit pairs.

To study the stability of cross-correlograms between FAM and NOV sessions, we inspected the between-session change in theta index, near-zero index, and symmetry score. There was a significant positive correlation between theta index of hippocampal pairs from QUINP rats in the FAM and NOV sessions (r=0.43, p<0.01; Fig. 5G). The correlations between theta index of hippocampal pairs in the CONTR group (r=-0.25, p=0.16) as well as ACC pairs in both groups (r=-0.11, p=0.83 for CONTR and r=0.02, p=0.94 for QUINP; Fig. 5H) were not significant. No significant difference between QUINP and CONTR was found when comparing the amount of inter-session change in theta index, neither for hippocampal (ranksum=1621, p=0.17) nor for ACC (ranksum=102, p=0.31, Wilcoxon rank-sum tests) unit pairs.

Near-zero index showed no significant correlation between the FAM and NOV sessions, neither for hippocampal (r=0.24, p=0.15 for CONTR and r=0.13, p=0.37 for QUINP group) nor for ACC (r=0.09, p=0.86 for CONTR and r=0.14, p=0.52 for QUINP group). We observed no significant difference in the inter-session change of near-zero index between the CONTR and QUINP groups (ranksum=1558, p=0.28 for hippocampal and ranksum=92, p=0.66 for ACC pairs, Wilcoxon rank-sum tests). There were no significant correlations between symmetry scores in the FAM and NOV session for either hippocampal pairs (r=0.27, p=0.11 for CONTR and r=-0.14, p=0.34 for QUINP group; Fig. 5I) or ACC pairs (r=-0.11, p=0.83 for CONTR and r=0.016, p=0.94 for QUINP group; Fig. 5J). There was no significant treatment effect on absolute value of the change of symmetry score between FAM and NOV session neither in the hippocampus (ranksum=1627, p=0.15) nor in the ACC (ranksum=94, p=0.58, Wilcoxon rank-sum tests).

Next, we assessed the dynamic structure of pairwise correlations within recording sessions at timescales of seconds. Each one-second interval of data was divided into 20 ms bins, and Pearson’s correlations of spike counts between a pair of units were calculated (See Methods). Change in correlations between consecutive intervals was measured as the mean absolute value of the difference in correlation coefficients. In the FAM session, pairwise correlations were changing significantly less in QUINP than in CONTR rats (ks=0.17, p<0.01; Fig. 5K). The same was true when measured over 5-second intervals, each divided into 100ms bins (ks=0.14, p<0.05). When inspecting longer intervals, there was no significant difference in pairwise correlation stability between QUINP and CONTR groups.

The stability of pairwise correlations between the FAM and NOV sessions was assessed next. For this analysis, the entire session was divided into 100ms bins. Pairwise correlation values of hippocampal pairs from both groups were significantly positively correlated between sessions, and the correlation was much higher for QUINP group (r=0.85, p<0.001) than for CONTR group (r=0.26, p<0.01; Fig. 5O). Hippocampal pairwise correlations have changed significantly less in the QUINP group (ranksum=16899, p<0.001, Wilcoxon rank-sum test). We also found significant correlation for ACC pairs in QUINP group (r=0.79, p<0.001) and in CONTR group (r=0.39, p<0.001; Fig. 5P). In the ACC, the inter-session change in pairwise correlation was significantly higher in the QUINP group (ranksum=453, p<0.001, Wilcoxon rank-sum test).

### Local field potential

Local field potential (LFP) activity was recorded from the hippocampal CA1 and ACC of quinpirole-treated (N=5) and control (N=5) rats. First, we inspected the overall frequency composition of LFP divided into two-second intervals using FFT (Fig. 6A1 and 6B1). As expected, the peak of power in the theta frequency range was more prominent in the hippocampal LFP compared to ACC in both groups (Fig. 6C and 6D). CONTR and QUINP groups did not significantly differ in relative theta power (t(8)=0.16, p=0.88 for the hippocampus and t(8)=-0.20, p=0.84 for the ACC; Fig. 6E).

**Figure 6.**
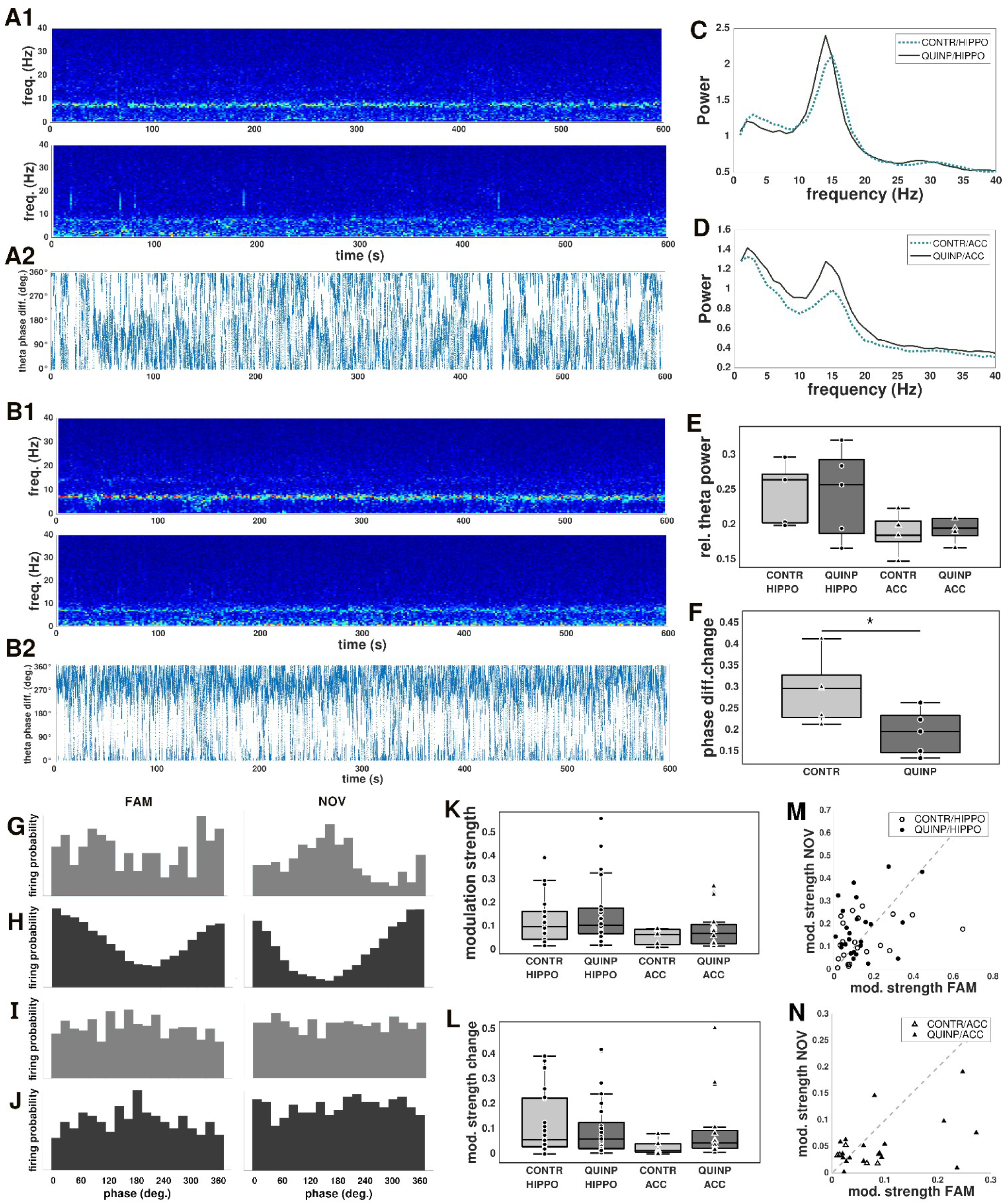
Dynamics of LFP oscillations – Experiment 2. **(A1)** An example of Fast-Fourier transformed LFP traces from the hippocampus (top) and ACC (bottom) of a CONTR rat in FAM session. **(A2)** An example of the phase difference between hippocampal and ACC theta oscillation from a CONTR rat. **(B)** Same as in (A), but for an example of a QUINP rat. **(C)** and **(D)** Mean hippocampal (C) and ACC (D) power spectra from CONTR (dashed line) and QUINP (solid line) rats in the FAM session. **(E)** Relative power in the theta frequencies in the FAM sessions. **(F)** Theta phase difference change between consecutive 20-s intervals. **(G)**, **(H)**, **(I),** and **(J)** Examples of theta phase locking of unit firing of the same four units in the FAM (left column) and NOV (right column) session. (G) Hippocampal unit from a CONTR rat. (H) Hippocampal unit from a QUINP rat. (I) ACC unit from CONTR rat. (J) ACC unit from QUINP rat. **(K)** Mean modulation strength of theta phase locking of units in the FAM session. **(L)** The change in mean modulation strength of theta phase locking between the first and the second half of the FAM session. **(M)** and **(N)** The modulation strength of theta phase-locking of units in the FAM session plotted against modulation strength in the NOV session for hippocampal (M) and ACC (N) units.

We calculated theta phase shift as the difference between momentary theta phase in the hippocampus and ACC (Fig. 6A2 and 6B2). To investigate the stability of theta phase shift within a recording session, we divided the LFP into 20-s long intervals. The intra-session variance of the theta phase shift among 20-second intervals, was significantly lower in QUINP rats when compared to CONTR rats (t(8)=2.32, p<0.05; Fig. 6F). The rate of change of theta phase difference in a session was also lower in the QUINP group, however the difference was not significant (t(8)=1.94, p=0.09). The variance of theta phase shift was also significantly lower in the QUINP group when 30-s intervals were analyzed. There was no significant difference when inspecting shorter (1s, 5s, and 10s) or longer (60s, 100s, 120s, or 300s) intervals.

We investigated the modulation of the firing of individual units by the theta rhythm. For each unit, we calculated the distribution of neuron’s probability to fire in different phases of local theta rhythm (Fig. 6G, 6H, 6I, and 6J). The magnitude of theta modulation of neuronal firing did not significantly differ between treatment groups (t(45)=0.04, p=0.97 for hippocampal, t(23)=-0.89, p=0.38 for ACC units; Fig. 6K). We measured the stability of the magnitude of theta modulation by the difference between the first and second half of the FAM session. There was no significant difference in how much theta modulation had changed in hippocampal units (ranksum=517, p=0.61; Fig. 6L) and ACC units (ranksum=43, p=0.06). Finally, we set out to study the stability of theta phase-locking between FAM and NOV sessions. There was a significant positive correlation between the mean magnitude of theta phase-locking during FAM and NOV sessions in hippocampal units from the QUINP group (r=0.57, p<0.01; Fig. 6M), but not in the control group (r=0.33, p=0.14). The correlation values for ACC units from QUINP rats (r=0.47, p=0.05) as well as CONTR rats (r=-0.52, p=0.29) were not significant (Fig. 6N). When comparing the inter-session change in theta phase locking between groups, we observed no significant difference, neither in the hippocampus (ranksum=507, p=0.96) nor in the ACC (ranksum=73, p=0.97, Wilcoxon rank-sum tests).

## DISCUSSION

We have studied the spatial and temporal dynamics of behavior and neuronal activity in a rat quinpirole-induced model of OCD. In summary, at the behavioral level, we observed increased locomotion and an increase in speed and acceleration within a session in the QUINP rats. The QUINP rats preferred repeated visits to the corners of the arena that contained objects. They also repeatedly visited the same sequences of locations, with increasing frequency of visits within a session. At the level of neuronal activity, we focused on the stability of neuronal firing patterns in space and in time. Quinpirole treatment was associated with decreased spatial information of the hippocampal units and decreased stability of spatial firing between sessions in ACC units. Quinpirole treatment led to increased stability of the organization of hippocampal neuronal firing in time. Theta modulation of hippocampal autocorrelograms showed greater stability within a recording session in the QUINP group. Similarly, the pairwise coactivation of units showed a tendency for increased inter-session and between-session stability in the QUINP group. Additionally, analysis of the phase difference between ACC and hippocampal theta also revealed greater stability in QUINP animals. Quinpirole effects on ACC units were more variable, showing both within-session increases in firing rate and less stable measurements between sessions.

Behavioral manifestations of the chronic quinpirole treatment that we observed confirm and extend earlier findings. Observed repeated visits to certain locations after quinpirole were analogous to reports of (Szechtman et al., 1998). Notably, in our study, the locations the rats returned to most often (corners with objects) were different from locations where they spent most time (corners without objects). In quinpirole-treated rats, we identified sequences of locations that were visited repeatedly and stereotypically in a ritual-like manner. These sequences were rat specific. We further noticed characteristic changes in behavioral dynamics within the duration of a session. In quinpirole-treated rats, average speed, acceleration, and stereotypy of locomotion increased with time of a session, while control animals decreased locomotion and exploratory behavior within an experimental session. Behavior in the two groups of animals did not significantly differ at the beginning of recording sessions, but increasingly diverged with increasing time in the arena. This extends reports of Janikova et al. (2019), who reported increased locomotion in quinpirole-treated rats within a session. While some of the quinpirole-induced changes, such as increased path length, speed, and acceleration, may reflect enhanced locomotion, the repeated spatial sequences capture more structured aspects of repetitive behavior beyond general hyperactivity.

The dynamic of increased frequency of stereotypical locomotory sequences with time is reminiscent of increasing discomfort during repeated checking behavior reported in human OCD. In humans, repeated checking paradoxically undermines confidence in memory and vividness of memory, although it does not affect memory accuracy. This phenomenon, observed in the general population (van den Hout and Kindt, 2003a, 2003b), is exacerbated in OCD patients (Maltby et al., 2005). Perhaps the gradually increasing frequency of stereotypical locomotion with time in rats may also be related to a gradual decrease in confidence in memory and increased discomfort.

Our observations reveal quinpirole-induced increased stability in the organization of neuronal activity in time, but no increased stability in the spatial organization of neuronal action potential discharge. This seemingly paradoxical observation is in line with previous reports of changes in the organization of hippocampal neuronal firing in time, often with preserved spatial tuning. Changes in the temporal structure of firing were observed, for example, after administration of NMDA antagonists that are used as psychotomimetic agents (Kao et al., 2017; Szczurowska et al., 2018) as well as after administration of cannabinoid receptor agonists (Robbe and Buzsáki, 2009). In our current observations and in the previous reports, changes in temporal organization of firing were associated with cognitive and behavioral alterations or impairments. These observations emphasize the need to look beyond “neural representations” and stimulus-response properties, and to focus also on the organization of “neuronal dynamics” as a key tool to understanding brain pathologies.

When considering neuronal dynamics changes associated with brain pathologies, concepts of attractor dynamics provide a particularly useful framework. In the context of OCD, this theoretical work posits that increased depth of attractor states that organize neuronal dynamics is the underlying cause of symptoms manifested as obsessive thoughts and compulsive actions (Rolls et al., 2008; Rolls, 2012). This theory predicts increased stability in firing patterns within neuronal networks in OCD pathology. We tested this prediction and our observation of increased stability of theta modulation of hippocampal firing, increased stability of correlations between neuronal activation, etc. provided supporting evidence for it.

Following quinpirole treatment, we observed an increased stability of theta modulation of single cell hippocampal activity within sessions. Theta rhythm has been implicated in several information-processing neuronal network functions in animal studies; it organizes activity of neuronal groups into cell assemblies (Harris et al., 2003), coordinates transitions between different representations (Jezek et al., 2011), and is crucial to modulation of learning-related plasticity in networks (Hölscher et al., 1997). Theta coordinates activity between the hippocampus and prefrontal areas, and this coordination is related to cognitive memory processes in rats (Jones and Wilson, 2005) as well as in humans (Su et al., 2024). Relationship between theta rhythm and neuronal network information flow in the cortico-hippocampal system was also shown in humans (Kragel et al., 2025). In human OCD, characteristic theta modulation of alpha power was observed during compulsions, and was interpreted as an indication of impaired coupling in cortical networks (Arbab et al., 2025). Against this background, our observations of increased stability of theta modulation of network activity after quinpirole treatment may have functional relevance for altered information processing and cognitive and behavioral impairments.

Beyond theta rhythm, at longer timescales of hundreds of milliseconds to seconds, we also observed higher stability of correlations in quinpirole-treated rats. At these timescales, dynamic changes in neuronal network activity related to attention, switching of representations, and memory retrieval have been characterized (Jackson and Redish, 2007; Fenton et al., 2010; Kelemen and Fenton, 2013). The specific roles of neuronal organization occurring at different timescales remain open to further exploration and promise novel insights into normal physiological function as well as pathological states.

The present study raises several important questions that could be addressed in future work. While we characterized behavior on the last day of quinpirole treatment, the activity of neuronal units was recorded for up to 10 days after the last dose of quinpirole. Einat and Szechtman (1993) reported that robust behavioral effects of chronic quinpirole administration using our protocol last up to 10 days after the last injection. In future research, recordings of neuronal activity performed in the same rats prior to and during the course of quinpirole administration could provide more detailed insights into changes in neuronal dynamics induced by this pharmacological procedure. Ensembles of large numbers of simultaneously recorded neurons in other cortical areas will allow analysis of aspects of network dynamics not accessible in the current work. Analyses extending beyond pairs of neurons, and focusing on the dynamics of large network states, may provide additional novel insights into neuronal information processing in pathologies like OCD.

To summarize, we characterized the dynamics of behavior in a quinpirole-induced model of OCD in rats. In a parallel experiment, we studied the dynamics of neuronal activity in the hippocampus and ACC in the same model. While the spatial organization of neuronal discharge was not substantially impaired in the quinpirole model, the temporal dynamics of neuronal firing displayed increased stability following quinpirole treatment, mostly in the hippocampus. We believe that these findings highlight the importance of studying neuronal network dynamics in pathological conditions to better understand the mechanisms underlying cognitive impairments.

## Declaration of generative AI use

The authors declare that they did not use generative AI to write this article. AI was consulted during programing the data analysis tools used in this project.

## CRediT authorship contribution statement

Adam F. Hanzlík: Data curation, Formal analysis, Investigation, Validation, Visualization, Writing - original draft, Software, Writing - review & editing. Ewa Szczurowska: Investigation, Methodology. Tereza Rydzyková: Investigation, Methodology. Eduard Kelemen: Conceptualization, Formal analysis, Funding acquisition, Methodology, Project administration, Software, Supervision, Writing - original draft, Writing - review & editing

## Ethical statement

This paper adheres to the principles for transparent reporting and scientific rigor of preclinical research as stated in Progress in Neuro-Psychopharmacology and Biological Psychiatry guidelines for Design and Analysis and as recommended by funding agencies, publishers, and other organizations engaged with supporting research.

## ACKNOWLEDGEMENTS

This work was supported by the Czech Science Foundation grants number 22-16717S and 26-23770S, both awarded to Karel Jezek and Eduard Kelemen, and by ERDF-Project Brain Dynamics, No. CZ.02.01.01/00/22_008/0004643. The authors thank Mia Anahi Ordonez for proofreading the manuscript.

## Notes

### Competing Interest Statement

The authors have declared no competing interest.

